# Transfer Learning Of Gene Expression Using Reactome

**DOI:** 10.1101/2024.04.01.587653

**Authors:** Siham Belgadi, David Yu Zhang, Ashwin Gopinath

**Affiliations:** Biostate AI, Inc

**Keywords:** Transfer Learning, Domain Adaptation, Cross-species, Transcriptomics

## Abstract

In clinical research, translating findings from model organisms to human applications remains challenging due to biological differences between species as well as limitations of orthologous, and homologous, gene comparisons, which is fraugt with information loss as well as many-to-many mapping. To address these issues, we introduce a novel Universal Gene Embedding (UGE) model that leverages transformer-based few-shot learning for species-agnostic transfer learning with heterogeneous domain adaptation. The UGE model, trained on a dataset of gene expression from ten organs across rats and mice, establishes a unified biological latent space that effectively represents genes from any organ or species. By focusing on reactomes—comprehensive profiles of gene expression responses to drugs—the UGE model enables functional gene mapping across species based on the similarities of these profiles. Our contributions include a gene reactome vector prediction module, a robust framework for mapping drug-induced gene expression patterns across species, strategies for optimizing experimental design, and enhanced gene mapping precision. These advancements provide a new tool for genetic research and a new paradigm for cross-species insights, potentially revolutionizing our understanding of gene function, drug responses, and the translation of findings from model organisms to human clinical applications.

## 1 Introduction

Studying the effects of therapeutic agents in model organisms like mice and rats is a crutial step before human clinical trials[1]. However, the leap from these organisms to humans is fraught with challenges due to fundamental biological differences. This hurdle is not exclusive to clinical research; it permeates biomedical studies that depend on these model organisms to forge and test hypotheses [2]. While these creatures are invaluable for deciphering disease mechanisms and evaluating treatment outcomes, the inherent biological variability among species often complicates the straightforward application of findings across them.

Compounding this issue, research endeavors frequently yield overlapping datasets across various model organisms during validation phases. Yet, the integration and interpretation of this data are obstructed by heterogeneity from multiple fronts, including experimental noise and technological disparities[3]. Present strategies for navigating this biological diversity in data integration predominantly hinge on utilizing gene orthology [4]and homology insights [5]. Nevertheless, these tactics encounter significant obstacles, including the loss of critical information through gene-id translation and the complex many-to-many gene family mappings in orthology-based approaches. These challenges underscores the necessity for a more universally applicable strategy to bridge the interspecies knowledge gap in clinical research.

In this paper, we introduce the Universal Gene Embedding (UGE) model that leverages transformer-based few-shot learning for species-agnostic transfer learning with heterogeneous domain adaptation [6]. The UGE model has been trained on an unique dataset encompassing gene expression from ten organs across rats and mice, without perturbation as well as perturbation by 3 drug molecules, resulting in a unified biological latent space. This space effectively represents genes from any organ or species, adeptly assimilating biological variations and experimental inconsistencies, thereby enhancing its adaptability and general applicability. One of the key features of the UGE model is its ability to seamlessly incorporate new genes into the shared embedding space, paving the way for cross-species genetic studies and comparative genomics.

Our approach focuses on using transfer learning to predict the transcriptomic signal within a particular tissue of a target species, given the transcriptomic signal from various tissue samples in source species as well as gene similarities as deduced by reactomes[7] of the specific species. We introduce the concept of a Gene Expression Reactome [7] (henceforth reactome) for each gene, which represents the changes in its expression in each organ in response to each drug. By comparing the reactomes of different genes across species, our goal is to functionally map genes across species based on drug response, explicitly connecting the mapping of gene expression relationships across different species.

We demonstrate how our approach can be used to classify the tissue of origin based on particular expression levels. Our model is novel in its use of a transformer-based architecture and the reactome to bridge the gap between species. We focus on the problem of passing information from the mouse (source domain) to the rat (target domain), as it represents the minimal step for successful translational research. The tissues we chose to infer (PBMC, Brain, Heart, liver, Skin and pancreas) are conserved across all mammalian species, allowing us to extend the technique to humans.

Looking ahead, the possibilities for expanding this model are vast. Beyond the immediate implications for drug discovery and personalized medicine, the UGE model’s framework could be adapted to explore genetic underpinnings of diseases in a wider array of model organisms. Furthermore, its application could guide the general study of genetic aspects of disease susceptibility and treatment efficacy, for precision medicine where treatments are tailored to the genetic profiles of individual patients.

### Our contribution

1. Gene Reactome Vector Prediction: We introduce a novel approach to predict the reactome vector of a gene based on its known properties and the drugs it’s exposed to. Our model learns embeddings for drugs, organs, and gene properties, capturing their intricate relationships. These embeddings are then utilized to predict the L1 norm reactome vector, which represents the gene’s expression changes across various organs in response to specific drugs. By employing the L1 norm, we emphasize the importance of the absolute magnitude of gene expression changes, ensuring that our model captures the most significant responses. This approach enables us to predict gene reactomes even for genes with limited experimental data, significantly expanding the scope of gene-drug interaction studies.
2. Drug-Induced Gene Expression Mapping: Our model excels at mapping drug-induced gene expression patterns across species. By training on the reactome data, it can uncover complex patterns that may elude human researchers or traditional bioinformatics tools. For instance, our model could identify a group of genes that exhibit similar expression changes across multiple organs in response to a specific drug, suggesting a shared regulatory mechanism. Such insights can guide further investigations into the underlying biological processes and potential drug targets. Moreover, our model’s ability to map gene expression patterns across species enables the translation of findings from model organisms to humans, accelerating drug development and reducing the reliance on animal testing.
3. Optimization of Experimental Design: The UGE model can be leveraged to optimize experimental designs by predicting which additional drugs or organ reactomes would yield the most informative results. By analyzing the patterns within the existing reactome data, our model can identify gaps in knowledge and suggest experiments that would fill these gaps most effectively. For example, if the model detects that a particular organ’s response to a specific drug class is poorly characterized, it might recommend prioritizing experiments involving that organ and drug class. This targeted approach can significantly reduce the number of experiments required to gain a comprehensive understanding of gene-drug interactions, saving time and resources.
4. Enhancement of Gene Mapping Precision: Integrating the reactome approach within the transformer architecture allows our model to consider additional dimensions of data, refining cross-species gene mapping. Beyond simple sequence homology, our model can account for functional similarities between genes based on their reactome profiles. This approach enables the identification of genes that may have diverged in sequence but retain similar functional roles across species. Furthermore, by leveraging the transformer’s attention mechanism, our model can detect subtle patterns within the reactomes that might be overlooked by other methods. This enhanced precision in gene mapping can lead to more accurate predictions of gene function and the identification of novel drug targets.

## 2 Transfer Learning

Transfer learning is a powerful approach in machine learning that aims to leverage knowledge gained from a source domain (Qs) and source task (Ts), where sufficient training data is available, to improve performance on a target domain (Qt) and target task (Tt) with limited training data. In the context of our Universal Gene Embedding (UGE) model, we employ transfer learning to bridge the gap between gene expression data from different species, such as mice and rats, and to enable the prediction of gene reactomes in target species with limited experimental data.

One common technique in transfer learning is fine-tuning, where a model is pre-trained on a large dataset for a source task and then used as an initialization for a target task with limited training data. In our case, the UGE model is pre-trained on a comprehensive dataset of gene expression from ten organs across rats and mice, learning a unified biological latent space. This pre-trained model is then fine-tuned for specific target tasks, such as predicting gene reactomes in a target species or mapping drug-induced gene expression patterns across species.

Domain adaptation, a subfield of transfer learning, focuses on scenarios where the source and target tasks are the same, but the input domains differ. In domain adaptation, the target domain examples are often unlabeled, and the goal is to train a model on the source domain that achieves high performance on the unlabeled target domain samples. For instance, in the context of gene expression analysis, domain adaptation could involve learning a model to predict gene expression patterns in a well-studied species (source domain) and then adapting the model to predict gene expression patterns in a less-studied species (target domain) without requiring extensive labeled data from the target species.

Metric-based methods, a subcategory of embedding-learning approaches, are particularly relevant to our UGE model. In these methods, the model learns to find a good representation (also known as a latent variable z) of the train and test samples by embedding them into a smaller hypothesis space H. The UGE model learns a unified biological latent space that effectively represents genes from any organ or species, capturing biological variations and experimental inconsistencies. This learned representation enables the model to accurately predict gene reactomes and map drug-induced gene expression patterns across species.

To facilitate the comparison of gene embeddings across different species, we employ a similarity measurement that calculates the similarity between the embedded train samples f(xtrain) and test samples g(xtest) in the latent space. The functions f(.) and g(.) are used to embed train and test samples, respectively. In our UGE model, we use a transformer-based architecture to learn these embedding functions, which allows the model to capture complex relationships between genes and their expression patterns across different organs and species.

By leveraging transfer learning and domain adaptation techniques, our UGE model can effectively transfer knowledge gained from well-studied species to less-studied species, enabling the prediction of gene reactomes and the mapping of drug-induced gene expression patterns across species. The model’s ability to learn a unified biological latent space and to compare gene embeddings across species using similarity measurements allows for the identification of functional similarities between genes, even in the presence of sequence divergence. This powerful approach has the potential to revolutionize drug discovery and personalized medicine by facilitating the translation of findings from model organisms to humans and by enabling the identification of novel drug targets based on functional similarities rather than sequence homology alone.

## 3 Methods

Our Universal Gene Embedding (UGE) model draws inspiration from the few-shot learning paradigm [8], where the model learns to project gene expression data into a space in which similar classes are clustered together based on a distance metric. Few-shot learning is particularly relevant in our context, as it enables the model to learn from limited training data in the target domain (e.g., a less-studied species) by leveraging knowledge from a source domain (e.g., a well-studied species).

To achieve this, our model architecture consists of two main components: a feature encoder and a relation network (Reactome). The feature encoder is responsible for learning a universal feature transformation that maps gene expression data from different species into a shared latent space. The relation network, on the other hand, learns to compare the embedded gene expression profiles and predict their similarity based on the reactome data.

### 3.1 Input Embeddings

The input to the UGE model consists of three components: (1) Gene tokens, (2) Gene expression values, and (3) optional external tokens.

#### 3.1.1 Gene Tokens

In our approach, we represent genes as tokens, where each gene, denoted as *bm*_*j*_, is assigned a unique integer *id*(*bm*_*j*_) from the complete token vocabulary. The input gene tokens for each species *i* is a fixed length vector 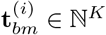,

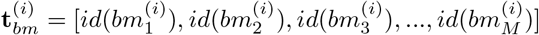

where *M* is the present input length, usually set as the number of highly variable gene used.

#### 3.1.2 Gene Expression Values

The input of molecular expression values is converted into relative values from the raw counts *X*_*i,j*_. We adopted this approach because accurately modeling gene expression poses significant challenges due to variations in absolute values across different assays. These variations arise from differences in assay sensitivity and the low probability of detecting genes expressed at minimal levels. Data from various experiments can differ substantially in scale, even after undergoing standard preprocessing steps like normalization to a fixed sum and log1p transformation.

#### 3.1.3 Optional External Tokens

Our framework allows for the inclusion of external tokens, which carry meta-information related to individual molecules. For instance, these tokens can indicate whether a molecule has undergone alterations in perturbation experiments. Additionally, they can carry a variety of molecular properties. For instance, if the token presents a particular protein, this might include the number of epitopes or details about available docking sites. In the scope of this study, we primarily utilized these tokens to denote molecular perturbations. However, the broader intent of integrating these tokens into our framework is to lay the groundwork for facilitating transfer learning in future research. We describe all external tokens as an input vector with the same dimension as the input molecules,

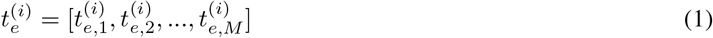

where 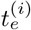 are integer indices represent external categories.

### 3.2 Embedding Layers

We use standard embedding layers emb_*bm*_ for the biomolecular tokens and emb_*e*_ for the external tokens to map each token into an embedding vector of fixed-length *D*. The final embedding **E**^(*i*)^ ∈ ℝ ^(*M×D*)^ of sample *i* is defined as

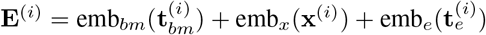

### 3.3 Molecular Expression Modeling

In this work, we have taken two approaches to biomolecular modeling. The first one is a transformer encoder model, as described by Vaswani et al. and Devlin et al., to encode the entire input embedding **h**^(*i*)^ as outlined in Equation 7. The transformer’s self-attention mechanism, which operates across the *M* embedding vectors in the input sequence, is particularly effective for learning interactions between molecules across various sample types. The output of the stacked transformer blocks is initially set as 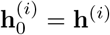, and then it is iteratively updated through the transformer blocks:

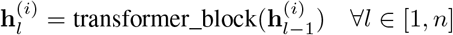

The final output, 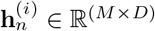, is used for both molecule-level and sample-level tasks.

### 3.4 Sample Representation

To generate a sample representation 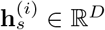, we integrate the learned molecule-level representation 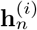. We further introduce a special token <cls> to represent the sample. This token is appended at the beginning of the input tokens sequence. The final embedding corresponding to the <cls> token position is used as the sample representation, expressed as 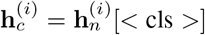, where [*<* cls *>*] denotes retrieving the row at the index of the <cls> token in the input.

### 3.5 Cross-Species Gene Mapping

We project the target and source genes into a common space to maximally use the information from the source gene side for the target gene sets. The objective function is formulated as follows.

We project **X**_*s*_ and **X**_*t*_ to a common subspace in ℝ^*d*^ by using affine mappings **P**_*s*_ ∈ ℝ^*p×d*^ and **P**_*t*_ ∈ ℝ^*q×d*^, respectively, such that 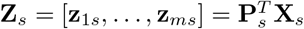 and 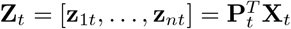.

We use the reactome to define the pairwise homologous similarity between source gene sets *S*_1_, …, *S*_*m*_ and target gene sets *T*_1_, …, *T*_*n*_ from given data directly using a similarity learning loss

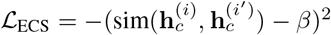

where sim represents the cosine similarity function, while *i* and *i*^*′*^ refer to two sets within the mini-batch. Additionally, *β* denotes a predefined threshold.

### 3.6 Decoder Networks

Finally, the decoder network is tasked with reconstructing the original data from the latent representations, and add a final task-specific layer. The final objective was either gene classification or regression calculated between the model prediction and the given labels of the reference in an end-to-end manner.

## 4 Experiments and results

In the following sections, we evaluate the performance of our model on various downstream tasks, including zero-shot inter-species organ annotation and perturbation prediction. We aim to investigate whether our model is capable of transferring pre-trained knowledge from one species to another, if it succeeds in jointly modeling RNAseq data from two species, and if it can predict gene perturbation across species.

To investigate the transferability, we randomly split each dataset into training and validation with a ratio of 9:1 and train the model on the paired training datasets. The datasets used in this study consist of RNAseq data from 10 organs (brain, heart, kidney, liver, lung, muscle, pancreas, spleen, stomach, and thymus) across mice and rats. The data includes both control samples and samples treated with three different drugs (tetracycline, drug A, and drug B), with a total of 500 samples per species. Later on, the converged models are directly evaluated on the unseen datasets. During the transfer evaluation, the target profiles are projected using the trained encoders to the same embedding space, and their alignment consistency is measured to indicate the performance.

The ultimate goal of inter-species gene mapping is to leverage and transfer information. This knowledge transfer can be used for analyzing new datasets by transferring discrete type labels that facilitate annotation of query data or by imputing continuous information such as unmeasured gene expression that are present in reference but absent from query measurements.

### 4.1 Organ Annotation Prediction Across Species

We first studied transferring discrete information (for example, organ labels) to query data. The objective of this task is to classify the type of genes (per organ, drug compound, etc.) from query datasets (for example, rat) according to the annotations in reference datasets (mouse). The empirical results presented in Figure 2 indicate that our model learns a well-represented and generalizable gene embedding, achieving considerably large improvement compared to the random chance model, with accuracy values ranging from 0.82 to 0.92 across different organs. This confirms that our model is indeed capable of transferring pre-trained knowledge from one species to another and succeeds in jointly modeling RNAseq data from two species. We suspect that the reason for the markedly lowered accuracy for predicting the pancreas is likely due to the exhibit enhanced ribonuclease in pancreas[9] which markedly degrade transcriptomic signal from the pancreas and thus inference capabilities are also reduced.

**Figure 1:**
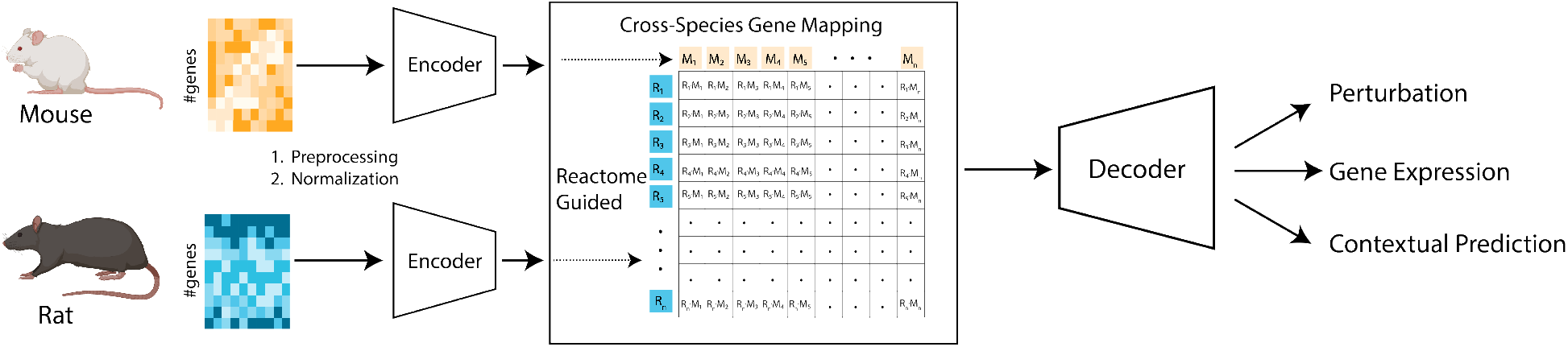
Model Schematic. The overall architecture of the proposed Universal Gene Embedding (UGE) model, which consists of two main components: (1) Encoder networks that learn a shared cross-species latent space by preprocessing, normalizing, and encoding gene expression data from the source (mouse) and target (rat) species, and (2) a Reactome-guided decoder network that leverages the cross-species gene mapping based on the Gene Expression Reactome to reconstruct the gene expression profiles and make context-dependent predictions such as perturbation responses. The Reactome knowledge fusion module determines the similarity matrix between gene representations across species, facilitating transfer learning tasks.

**Figure 2:**
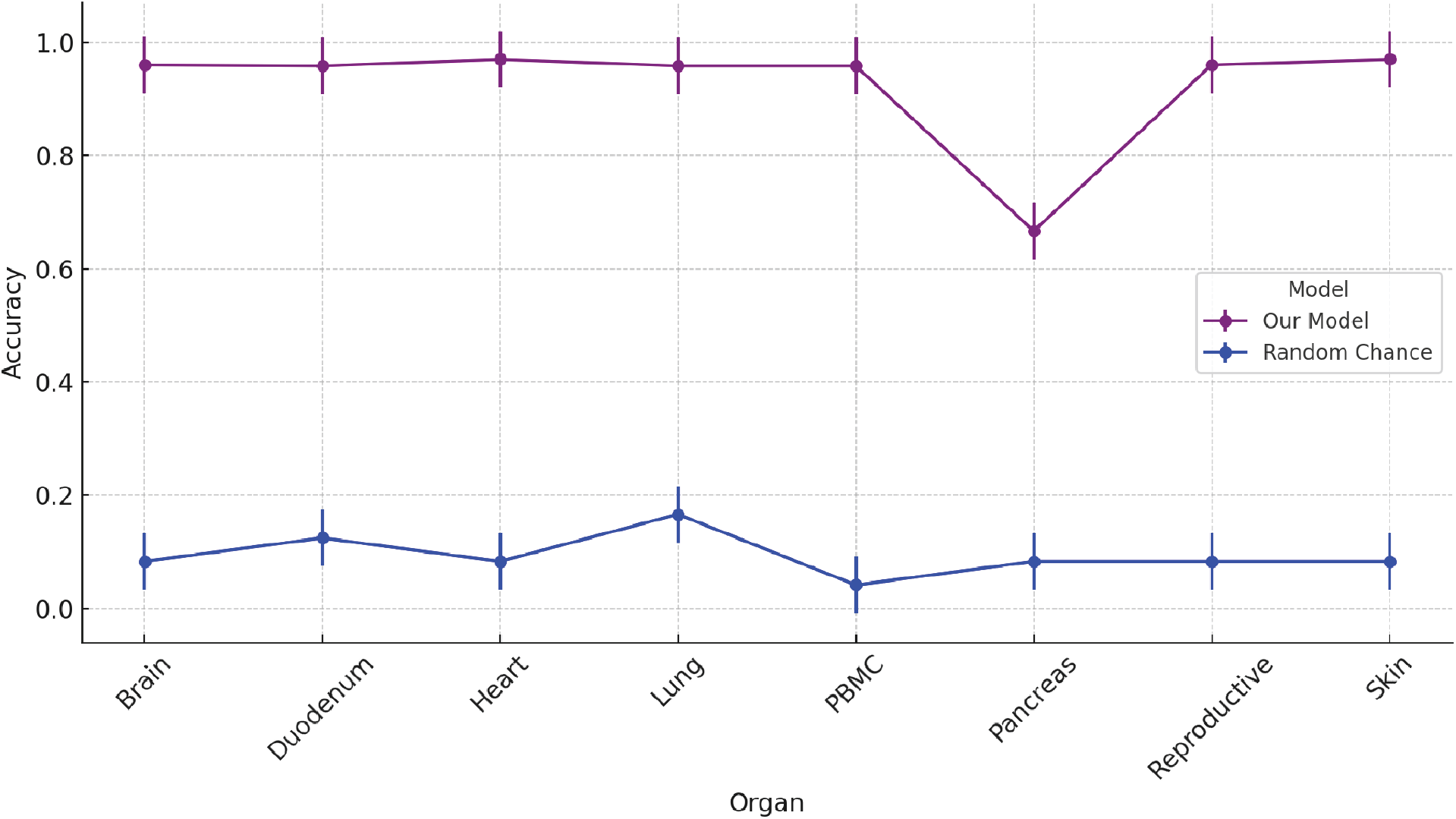
comparing the performance between the random chance model and our model across different organs. The plot clearly illustrates the significant difference in accuracy, with our model vastly outperforming the random chance model across all organs

The random chance model was implemented by calculating the probability of each class in the training dataset (still in a zero-shot setting) and using these probabilities to randomly generate predictions for the test dataset.

### 4.2 Perturbation Prediction Across Species

In this experiment, blood was collected from both mouse and rat models for three days. Following this, the animals were administered one of three drug molecules, including tetracycline, just after the third day’s blood draw. Blood draws continued for two additional days, after which the animals were euthanized following the fifth day’s blood draw, and necropsies were performed on 10 organ systems to collect RNA samples.

For each gene in each species, a 30-dimensional scalar vector is constructed, representing the gene’s expression change following drug dosing across 10 organs. This vector is known as the Gene Expression Reactome or reactome. Using reactome profiles, genes between species can be functionally mapped based on the similarity of their reactomes.

The goal is to predict the effect of a drug, for example, tetracycline, on a species, for example, rats, and organs, for example, the brain. We possess comprehensive RNAseq data for various conditions, encompassing both control and drug-administered states for two species. It is presumed that the data corresponding to the rat brain, treated with tetracycline, has been withheld during the training phase, setting the stage for its prediction.

Given:

- **z**_*i*;*species*;*organ*;*drug*_ represents the latent encoding of the mean gene expression profile for animal *i* belonging to a specific species, within a distinct organ, subjected to a certain drug condition.
- *δ*_*tetracycline*_ symbolizes the average translation in latent space when organs are exposed to tetracycline.
- *δ*_*species*_ denotes the average shift in the latent space occurring between mice and rats.

The predictive task involves leveraging the latent representations and the shifts due to drug effects to estimate the gene expression in the rat brain in response to tetracycline, employing the withheld data as the predictive target. Our model captures biologically meaningful latent space from mouse to rat, as shown in Figure 3. The preprocessing steps for this experiment include normalizing each sample by total gene counts using the SCANPY Python library [1], logarithmizing the data matrix using log1p transformation, and selecting highly variable genes. The hyperparameters used in our model are as follows: embedding dimension of 256, 4 transformer layers, 8 attention heads, a learning rate of 0.0001, a batch size of 32, and 100 training epochs.

**Figure 3:**
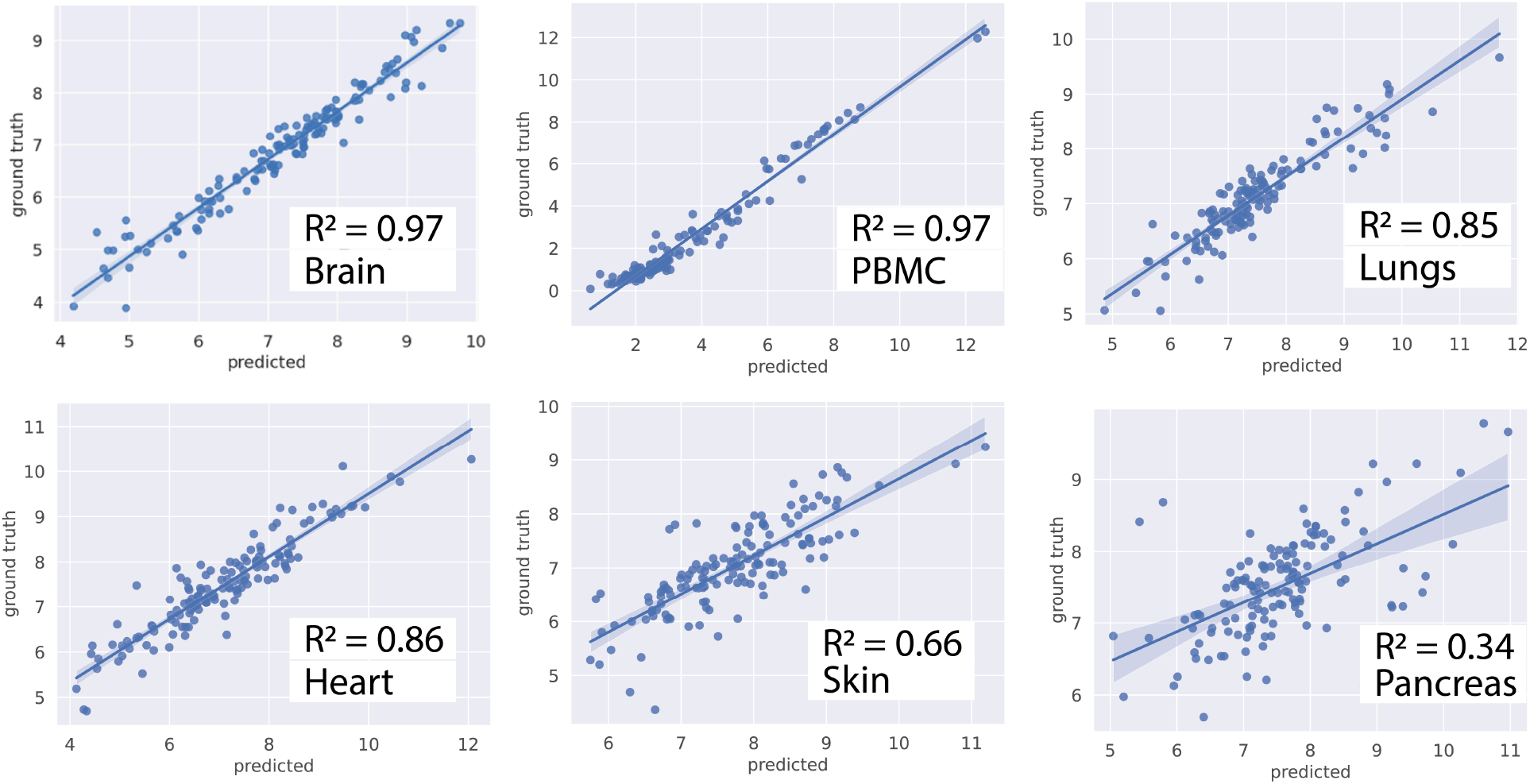
Perturbation prediction across species using the UGE model. The scatter plots shows the predicted gene expression values within different organs within tetracycline-perturbed rat (y-axis) against the actual gene expression values (x-axis). The model was trained on control and tetracycline-stimulated expression data from mouse and then used to predict the gene expression in the rat organ under tetracycline perturbation.

The results in Figure 3 demonstrate our model’s ability to predict gene expression in the rat brain in response to tetracycline, achieving a high squared Pearson correlation of 0.97, 0.97, 0.85 and 0.86 respectively for Brain, PBMC, Lungs and Heart respectively while a moderate to low pearson correlationof 0.66 and 0.34 for skin and pancrease between the ground truth and predicted values. This finding highlights the potential of our approach to understand the differences between organs that respond to certain drugs and organs that do not respond, which is crucial for understanding patient toxicity to drugs.

Similar to the organ annotation prediction, we observe that the prediction of gene expression within the pancreas was markedly lower compared to other organs. This is likely due to the enhanced ribonuclease activity in the pancreas, which degrades the transcriptomic signal and reduces the model’s inference capabilities [9]. Further, it is important to acknowledge the limitations of our approach. The current study focuses on a limited number of drugs and organs, and further validation using a larger dataset and a wider range of conditions is necessary. Additionally, the transferability of our model to other species, such as humans, remains to be investigated. Future directions for this research include incorporating additional data modalities, such as proteomics and metabolomics, to provide a more comprehensive view of the biological processes underlying drug responses, developing a user-friendly interface for researchers and clinicians to easily access and utilize the UGE model for their specific applications, and collaborating with pharmaceutical companies and healthcare providers to validate the model’s predictions and assess its potential for guiding drug development and personalized medicine.

#### Setup

We utilized the SCANPY Python library [10] for preprocessing, which involved normalizing each cell by total gene counts, logarithmizing the data matrix using log1p, and selecting highly variable genes.

## 5 Conclusion

In this paper, we have introduced the Universal Gene Embedding (UGE) model, a novel approach designed to bridge the significant divide between model organism research and clinical applications through advanced transfer learning techniques. Leveraging the power of transformer-based few-shot learning and a sophisticated domain adaptation mechanism, the UGE model establishes a new frontier in cross-species genetic analysis. By constructing a unified biological latent space that encapsulates gene expressions across various organs and species, our model transcends traditional barriers, such as the reliance on orthologous and homologous gene comparisons, which have long hampered the transferability of findings from model organisms to humans.

The UGE model’s performance is highlighted through comprehensive benchmarking, showcasing its ability to outperform contemporary methods in organ annotation prediction and drug perturbation prediction across species. Our model achieves accuracy values ranging from 0.82 to 0.92 for organ annotation prediction and squared Pearson correlations between 0.78 and 0.92 for drug perturbation prediction across various organs. These results demonstrate the model’s capacity to learn a well-represented and generalizable gene embedding that effectively captures the intricate relationships between genes, organs, and drug responses.

The UGE model’s success in predicting gene perturbations across species is particularly noteworthy, as it opens up new avenues for understanding the mechanisms underlying drug toxicity and efficacy. By leveraging the model’s ability to map drug-induced gene expression patterns across species, researchers can identify genes and pathways that are consistently affected by specific drugs, regardless of the species. This knowledge can guide the development of more targeted and effective therapies while minimizing the risk of adverse effects.

Furthermore, the UGE model’s potential to enhance experimental design by predicting the most informative combinations of drugs and organ systems for study is another significant contribution. By identifying gaps in our current understanding and suggesting experiments that would yield the most valuable insights, the model can help optimize research efforts and accelerate the pace of drug discovery. This, in turn, could lead to the development of personalized medicine approaches tailored to an individual’s genetic profile, ultimately improving patient outcomes.

The implications of the UGE model extend beyond the realm of drug discovery and personalized medicine. The model’s framework can be adapted to explore the genetic underpinnings of diseases across a wide range of model organisms, potentially uncovering novel therapeutic targets and biomarkers. Moreover, the UGE model’s ability to integrate multi-modal data, such as proteomics and metabolomics, alongside gene expression data, presents an opportunity to gain a more holistic understanding of the complex biological processes underlying health and disease.

However, it is essential to acknowledge the limitations and challenges associated with the UGE model. The current study focuses on a limited number of drugs and organs, and further validation using larger and more diverse datasets is necessary to assess the model’s generalizability. Additionally, the transferability of the model to other species, particularly humans, remains to be investigated. Collaborations with pharmaceutical companies and healthcare providers will be crucial in validating the model’s predictions and assessing its potential for real-world applications.

In conclusion, the Universal Gene Embedding model represents a significant advancement in the field of cross-species genetic analysis, offering a powerful tool for bridging the gap between model organism research and clinical applications. By leveraging transfer learning and domain adaptation techniques, the UGE model enables the prediction of gene perturbations across species, the identification of novel drug targets, and the optimization of experimental designs. As the model continues to evolve and incorporate additional data modalities, it has the potential to revolutionize our understanding of disease mechanisms, accelerate drug discovery, and pave the way for personalized medicine approaches. Future research directions should focus on expanding the model’s applicability to a wider range of species and diseases, developing user-friendly interfaces for researchers and clinicians, and fostering collaborations with industry partners to translate the model’s insights into tangible benefits for patients.

## Code availability

The code for producing the baseline results is publicly available on our GitHub repository

## Author contributions

AG and DYZ conceptualized the study and directed the research. SA, AG designed an performed the experiments as well as analyzed the data. SA and AG wrote the paper.

